# Revisiting the Role of Notch in Nephron Segmentation: Notch is Required for Proximal, Not Distal Fate Selection During Mouse and Human Nephrogenesis

**DOI:** 10.1101/2021.12.14.472634

**Authors:** Kathryn Duvall, Lauren Bice, Alison J. Perl, Naomi Pode Shakked, Praneet Chaturvedi, Raphael Kopan

**Affiliations:** Department of Pediatrics, University of Cincinnati College of Medicine, Cincinnati, Ohio, USA; Division of Developmental Biology, Cincinnati Children’s Hospital Medical Center, Cincinnati, Ohio, USA; Division of Nephrology and Hypertension, Cincinnati Children’s Hospital Medical Center, Cincinnati, Ohio, USA; Sackler Faculty of Medicine, Tel Aviv University, Israel

## Abstract

Notch signaling promotes maturation of nephron epithelia, but its proposed contribution to nephron segmentation into proximal and distal domains has been called into doubt. We leveraged single cell and bulk RNA-seq, quantitative immunofluorescent lineage/fate tracing, and genetically modified human iPSC to revisit this question in developing mouse kidneys and human kidney organoids. We confirmed that Notch signaling is needed for maturation of all nephron lineages, and thus mature lineage markers fail to detect a fate bias. By contrast, early markers identified a distal fate bias in cells lacking *Notch2*, and a concomitant increase in early proximal and podocyte fates in cells expressing hyperactive *Notch1* was observed. Orthogonal support for a conserved role for Notch signaling in the distal/proximal axis segmentation is provided by the ability of *Nicastrin*-deficient hiPSCs-derived organoids to differentiate into TFA2B+ distal tubule and CDH1 connecting segment progenitors, but not into HNF4A+ or LTL+ proximal progenitors.

**Summary:** Notch signaling acts in nephron segmentation to select early proximal, but not distal tubule fate downstream of a global role promoting epithelial growth and maturation in mouse and human.

## Introduction

The evolutionarily conserved Notch pathway translates an *inter*-cellular interaction into *intra*-cellular transcriptional outputs that control cell fate, proliferation, differentiation, and apoptosis in a context-specific manner (Artavanis-Tsakonas et al., 1999; Bray, 2006; Kopan and Ilagan, 2009; Kovall et al., 2017). Mammals possess four Notch receptors (N1 to N4) and five Delta/Jagged ligands; all produce a signal when a bound ligand, presented and endocytosed by a neighboring cell, applies force that unfolds the Notch juxtamembrane negative regulatory region, enabling cleavage by the metalloprotease ADAM10. The truncated, cell membrane bound polypeptide is then cleaved by the γ-secretase complex freeing the <u>N</u>otch <u>I</u>ntra<u>c</u>ellular <u>D</u>omain (NICD), which translocates into the nucleus (Kopan and Ilagan, 2009; Kovall et al., 2017). NICD associates with the DNA-binding protein CSL (<u>C</u>BF1/<u>S</u>uppressor of Hairless/<u>L</u>AG-1, also known as RBPJ in vertebrates) and recruits the coactivator Mastermind to activate Notch target gene expression (Bray, 2016; Gordon et al., 2008; Kovall et al., 2017). Notably, each Notch receptor can be used only once, with NICD degrading after association with the transcription machinery (Kuang et al., 2020).

Mammalian kidneys develop from a Six2-expressing progenitor population induced by the ureteric bud (UB) to transition from self-renewal to differentiation via mesenchymal-to-epithelial transition (MET), forming a multitude of nephrons (Costantini and Kopan, 2010). While there is broad agreement that the Notch pathway plays an important role in nephron formation (nephrogenesis), some controversy exists as to the specifics of its contributions to this process in mammals. After MET, cells form a pre-tubular aggregate (PTA) and start proliferating, forming a renal vesicle (RV) which contorts into the S-shaped body (SSB), where it is thought future nephron linages are already set (Lindstrom et al., 2018; Lindstrom et al., 2021). Early reports using inhibitors of γ-secretase (Cheng et al., 2003) or hypomorphic PSEN1 transgene in γ-secretase null mutants (Wang et al., 2003) suggested that Notch signals are required in a temporal window starting after MET and ending after formation of the SSB, noting the paucity of nephron epithelia in these kidneys. In follow up studies, it was established that loss of a conditional Notch1 allele was tolerated, but removal of Notch2 from the posterior mesoderm (with Pax3Cre, (Cheng et al., 2007)) resulted in lethality due to paucity of nephrons. Some Wnt4-positive RV formed in these animals, but the transition to SSB failed and they developed no further. Cdh1 expression in *<u>L</u>otus <u>t</u>etragonolobus* <u>l</u>ectin (LTL), Wt1 and Krt8 triple-negative epithelia resembling an incomplete SSB was used to argue that the few surviving Notch2-deficient epithelial cells were distal in character. The reason for the limited epithelial endowment in Notch2 nulls was not explored beyond the demonstration that proliferation was reduced in Jag1+, Notch2 deficient cells relative to Jag1+ wild type cells. The assertion that Notch activity biased a distal default program towards a proximal tubule (PT) fate was further supported by the observation that activating a stabilized (degron-deleted) version of the Notch1 intracellular domain (N1ICD), the active form of the receptor, resulted in *en masse* differentiation of nephron progenitor cells (NPC) into multiple LTL or Wt1 positive proximal epithelial clusters lacking distal tubule or loop of Henle (LOH) markers (based on or explant cultures (Cheng et al., 2007) or RNA *in-situ* hybridization at embryonic day 15 (E15) (Boyle et al., 2011)).

Independent analysis of an activated, degron-containing N2ICD transgene (Fujimura et al., 2010)) arrived at a somewhat different conclusion. While they too observed abundance of proximal tubules and aborted branching consistent with precocious NPC differentiation, they also report a greater degree of abnormal differentiation with tubular and glomerular cysts. Distal markers were not examined. The glomeruli were far less disrupted than those over-expressing the stabilized N1ICD^ΔPEST^ (Sweetwyne et al., 2015; Waters et al., 2008). Notably, they concluded that Notch signals were required for epithelial maturation, not fate selection, and suggested that because Notch1 and Notch2 promote different outcomes, they may play different roles. The similarity of the NICD over expression phenotypes to that observed in Six2 null mutants, which undergo precocious nephron differentiation, was used to suggest that NICD inhibited Six2, led them to propose that Notch target genes may contribute to epithelization both indirectly (by repressing the NPC main maintenance factor, Six2) and directly (by promoting maturation).

The idea that Notch1 and Notch2 play distinct developmental roles due to divergent residues in their respective intracellular domains was tested *in vivo* directly by swapping the N1ICD with the N2ICD and vice versa (Liu et al., 2013). Pax3Cre-mediated deletion of *Notch2^f/f^* could not be rescued by substituting N2ICD into Notch1. By contrast, mice without N2ICD (now having 4 copies of Notch1 NICD, two replacing N2ICD) displayed normal kidney development. Because each Notch receptor is consumed as it generates a signal and cannot be reused, and because Notch2/Jag1 interactions produced more NICD polypeptides than Notch1/Jag1 interactions, it was concluded that signal strength, defined as the sum of NICD released from all ligand-bound Notch receptors on the cell surface, was a far more important determinant of Notch signaling outcomes in the kidney than NICD composition (Liu et al., 2015; Liu et al., 2013; Ong et al., 2006), providing a different interpretation for the differences between N1ICD^ΔPEST^ (strong) and N2ICD (weaker) over expression.

Several mechanisms have been found to modulate Notch1 vs Notch2 signal strength, including receptor glycosylation (Haltiwanger, 2008; Kakuda and Haltiwanger, 2017) which generates the preferred Notch2/Jag1 and Notch1/Dll1 signaling pairs, consistent with the more profound impact of deleting *Jag1* than *Dll1* (Liu et al., 2013). More recently, ligand-dependent signal dynamics were shown to be another key determinant of signaling outcomes (Nandagopal et al., 2018) with Dll1, and thus Notch1, signaling in a pulsatile pattern, whereas Jag1 and thus Notch2 produced a sustained signal. Collectively, these observations are consistent with the idea that the different outcomes described above reflect differences in the relative strength of NICD signals, be it endogenous or over expressed.

Whereas support for the role of Notch signaling in epithelial maturation has strengthened, a role for Notch signals in promoting proximal fates over distal ones has been called into doubt by these findings and (Chung et al., 2017). MET can be completed without Notch1/2 or RBP, but these kidneys have even fewer epithelia (Chung et al., 2016; Surendran et al., 2010a) than in mice with *Notch2* null nephron progenitor cells (NPCs) (Cheng et al., 2007) supporting a key role for the Notch pathway in transitioning from an epithelial vesicle to a nephron. Moreover, because the commonly used *Six2*^+*/Cre*^ line is mosaic, allowing some NPC to “escape” deletion altogether or delay it sufficiently, formation of epithelia in the Surendran study could reflect such escapers. Another confounder of earlier findings is the use of *Pax3Cre*, which deletes *Notch2* in both epithelia and mesenchyme. To address some of these issues, investigators examined differentiation markers for podocytes (*Nphs2*), proximal tubules (*Slc34a1*), distal tubules (*Slc12a3*), and LOH (*Slc12a1*) in mice deleting both *Notch1* and *Notch2* with *Wnt4*^+*/Cre*^, and reported that no recognizable epithelial linage formed from the SSB (Chung et al., 2017). Moreover, when N1ICD^ΔPEST^ was activated in nascent epithelia by Wnt4Cre, a cell fate bias was not observed based on these markers. The authors concluded that Notch signals have no role in segmenting the Proximo-distal axis in nephrons, but did not consider the possibility that since Notch is required for maturation, the absence of these late markers in loss of function (LOF) models may not be informative.

Among the relevant recent advances is the explosion of scRNA-seq studies, leading to the appreciation that first-generation markers, including the ones used by (Chung et al., 2017), describe relatively mature cell populations. A far more granular view of nephrogenesis emerged, with markers identifying intermediate steps towards formation of the adult nephron cell types (Naganuma et al., 2020). Using our own scRNA-seq data we positioned Notch receptor-ligand pairs in a temporal context along the differentiation axis, and revisited the original *Notch2* LOF model with new tools. We demonstrate herein that deletion of *Notch2* with *Six2C*^+*/Cre*^ or with *Fgf20*^+*/Cre*^, which in the kidney generate exclusive deletion in the NPC and their epithelial descendants, sufficed to arrest epithelial development at an immature state, indistinguishable morphologically from our earlier report (Cheng et al., 2007). Notch2 antibody staining and lineage analysis confirmed that the epithelial cells that formed were indeed descendants of *Notch2* deficient cells. Most *Notch2*-deficient, GFP-positive descendants of *Six2*^+*/Cre*^; *Notch2^f/f^*; *ROSA*^+*/EYFP*^ NPCs expressed *Cdh1* and *Tfap2b*, an early marker for committing distal tubules/LOH (Marneros, 2020; Naganuma et al., 2020). A small minority of GFP+ cells were positive for LTL and Hnf4a, a committing proximal tubule marker (Marable et al., 2020), but not for *Cdh1*. The inverse bias favoring the proximal fate was seen in *Six2*^+*/Cre*^; *N1ICD^ΔPEST^* kidneys, with abundant expression of Hnf4a+ cells and Wt1+ cells, but lacking Tfap2b+ cells. A handful of epithelial cells expressed all lineage markers (LTL, Hnf4a, Wt1, and Tfap2b). Similarly, *Six2*^+*/Cre*^; *RBP^f/f^* kidneys had far more Tfap2b+ cells than LTL/Hnf4a+ cells. Delayed deletion of *RBP^f/f^* with Wnt4^+/Cre^ did not compromise nephron formation, marking the end of the Notch-dependent window as the time required for RBP protein to turn over after the gene is deleted, with a half-life measured at 60 hours in the epidermis (Turkoz et al., 2016). At the transcriptome level, RNA-seq analysis confirmed a shift in gene expression towards early PT (ePT), and away from NPC, mature PT, DT or LOH in N1ICD^ΔPEST^-expressing NPC. Finally, human kidney organoids derived from *Nicastrin (NCSTN)* deficient hiPSC which have lost γ-secretase activity and thus, all Notch signals, differentiated into KRT8/18 and TFAP2B positive cells, but not into HNF4A or LTL positive cells. Combined, these analyses confirm a role for Notch signaling in proximal/distal nephron segmentation downstream of a more global role in promoting epithelial growth and maturation in both mouse and human.

## Results

### scRNA-seq places the Notch signaling pathway in early epithelia with a slight proximal distribution bias

As introduced above, the present understanding of Notch signals to nephrogenesis assumes that Notch signaling did not contribute to the epithelial fate selection process, but this conclusion was based on analyses using markers for mature cell fates. Single cell RNA sequencing (scRNA-seq) suggested that the choice of markers confounded the interpretation. Integration analysis of six cortically-biased scRNA-Seq datasets (pooled embryos from three genotypes with high transcriptional similarity, including E14 and P0 timepoints) using the Seurat 3.0 UMAP algorithm (Stuart et al., 2019) identified several super clusters (“continents”), characterized as NPC, stromal, and ureteric bud (UB, *Ret*+) continents (Jarmas et al., 2021). The NPCs were aligned in a continuum with (*Cited1*+, *Six2*+, *Wt1*+, *Nr2f2*+, *Pax8*+) naïve cells at one pole, followed by a transitional (*Cited1*−, *Six2*+, *Wt1*+, *Nr2f2*+, *Wnt4*+, *Pax8*+, *Lhx1*+) population (marked with red brackets), and segregating into epithelial renal vesicle and precursors of distal tubule (DT, *Fgf8*+, *Sim1*+, *Tfap2b*+, *Gata3*+), proximal tubule (PT, *Cdh6*+, *Hnf4a*+, *Fut4*+), and podocyte (Pod, *Mafb*+) at the opposite pole (Figure 1). Importantly, within each emerging epithelial cluster, the UMAP algorithm arranged cells from the least mature to the more mature, with the most mature marker (*Slc12A3* (DT), *Slc34a1* (PT) and *Nphs2* (POD) absent from this cortically biased dataset (Figure 1). Interrogating the expression of Notch pathway ligands (*Jag1*, *Dll1*), receptors (*Notch1*, *Notch2*) and targets (*Nrarp*, *Hes5*, *Hnf1b*, *Hey1*, *Hey2*, *HeyL*) shows expression in the transition zone and a strong distribution bias towards the proximal lineage, with minimal overlap with the *Tfap2b* domain (a marker for distal fates, Figure 1). This pattern rekindled our interest in clarifying the role of Notch in nephron segmentation.

**Figure 1:**
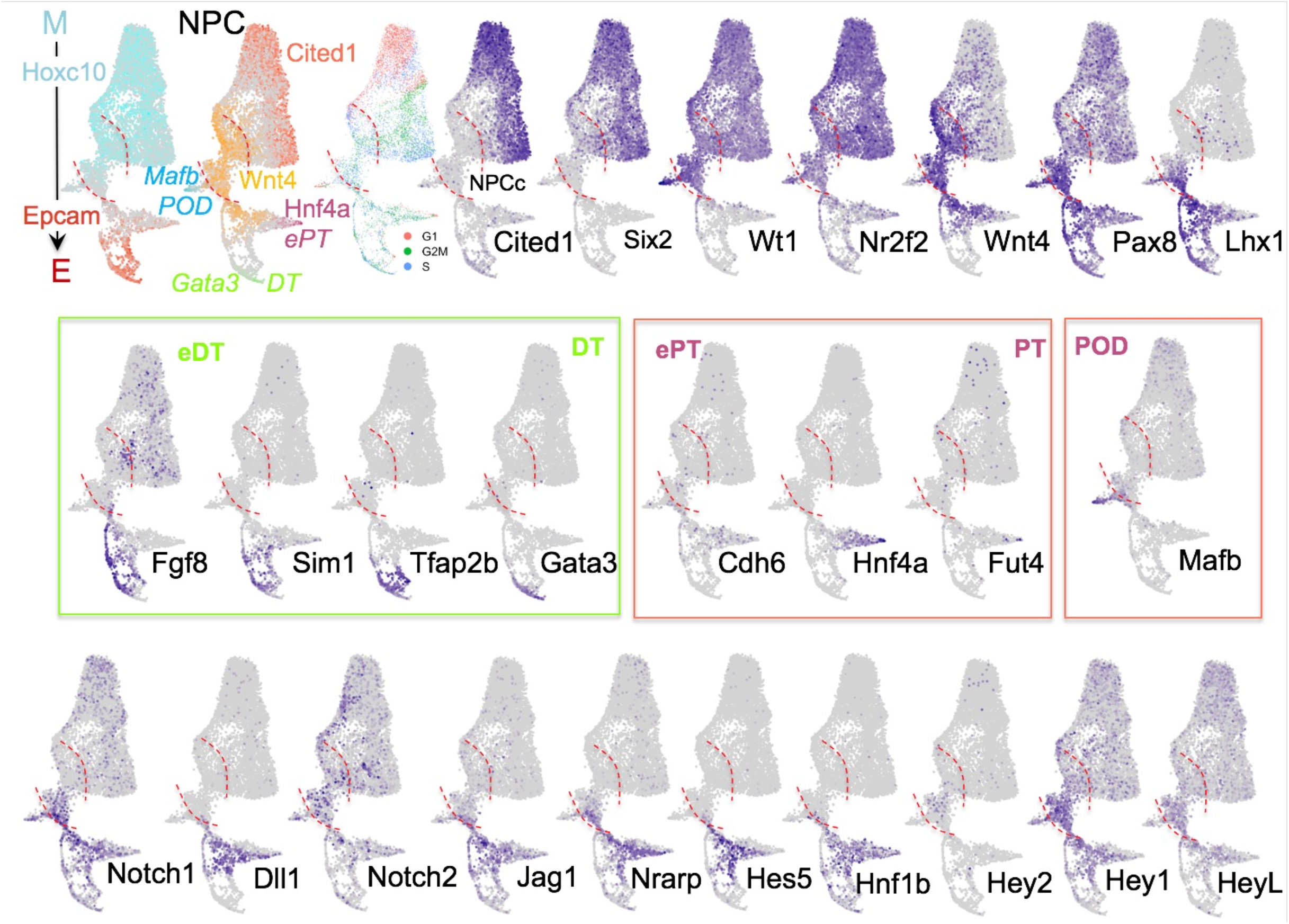
scRNA places the Notch pathway at PT/DT fate branching. Single cell RNA integration analysis of data collected from pool of E14 and P0 embryo from Six2-cre, Six2 KI, and Six2-cre; Tsc1^+/f^ (Jarmas et al., 2021). Top row, left orients reader to the mesenchymal (HoxC10) to epithelial (Epcam) axis, and marks cell population by colorized marker gens within the NPC cluster. Cell cycle distribution is also shown, with G1 cells mostly in the naïve NPC region (Cited1+). Red dotted parentheses indicate the transition zone (Wnt4+) from committed NPC (NPCc) to DT (Epcam+, Gata3+), ePT (Hnf4a+). In all panels. Area above the dashed red bracket contains the NPC (Cited1+, Six2+). Gene expression pattern are set such that all expressing cells are in front of non-expressing cells. Below the lower red bracket cells are transitioning or already epithelia. In the middle row, early marker for PT (ePT), DT (eDT), and POD. The bottom row depicts cells expressing Notch signaling components and targets. The Notch1/Dll1 and Notch2/Jag1 are preferred receptor/ligand pairs. Note, notch target genes may respond also to other stimuli. Nrarp is expressed most widely in Notch-responsive tissues and tumors.

### Notch2 Deletion only in NPC is sufficient to ablate nephrons and compromise viability

We were aided in our analysis by the unexplained change in the behavior of the widely used *Six2*^+*/Cre*^ strain upon establishment of the colony in a new environment. After establishing our mouse colony at Cincinnati Children’s Hospital Medical Center (CCHMC), we noted a change in the outcome of a cross between *Six2*^+*/Cre*^*; Notch2^+/f^* animals. In contrast to the outcome of this cross at Washington University School of Medicine (WUSM) (Surendran et al., 2010a), no viable *Six2*^+*/Cre*^*; Notch2^f/f^* offspring were obtained at CCHMC. Survival analysis on 10 litters (73 pups) confirmed that while *Six2*^+*/Cre*^*; Notch2^f/f^* pups were born at the expected Mendelian ratios, all perished by postnatal day 2 (*p*<10^-5^; Figure 2A), exactly as reported for the *Pax3*^+*/Cre*^*; Notch2^f/f^* pups. The complete penetrance of the phenotype argued against the presence of an unlinked lethal allele, and using multiple unrelated *Notch2^f/f^* dams with no surviving *Six2*^+*/Cre*^*; Notch2^f/f^* pups argued against a linked mutation at the *Notch2* locus. To rule out the possibility that our Six2TGC breading stock acquired a linked mutation segregating with Six2-Cre, we obtained a stud from Dr. Joo-Soep Park at CCHMC and again, all *Six2*^+*/Cre*^*; Notch2^f/f^* pups perished by P2. This result demonstrated that *Notch2* deletion only in NPC sufficed to reproduce the observations made by (Chung et al., 2017), eliminating the concern that *Notch2* loss in mesenchyme contribute to the reported phenotypes.

**Figure 2:**
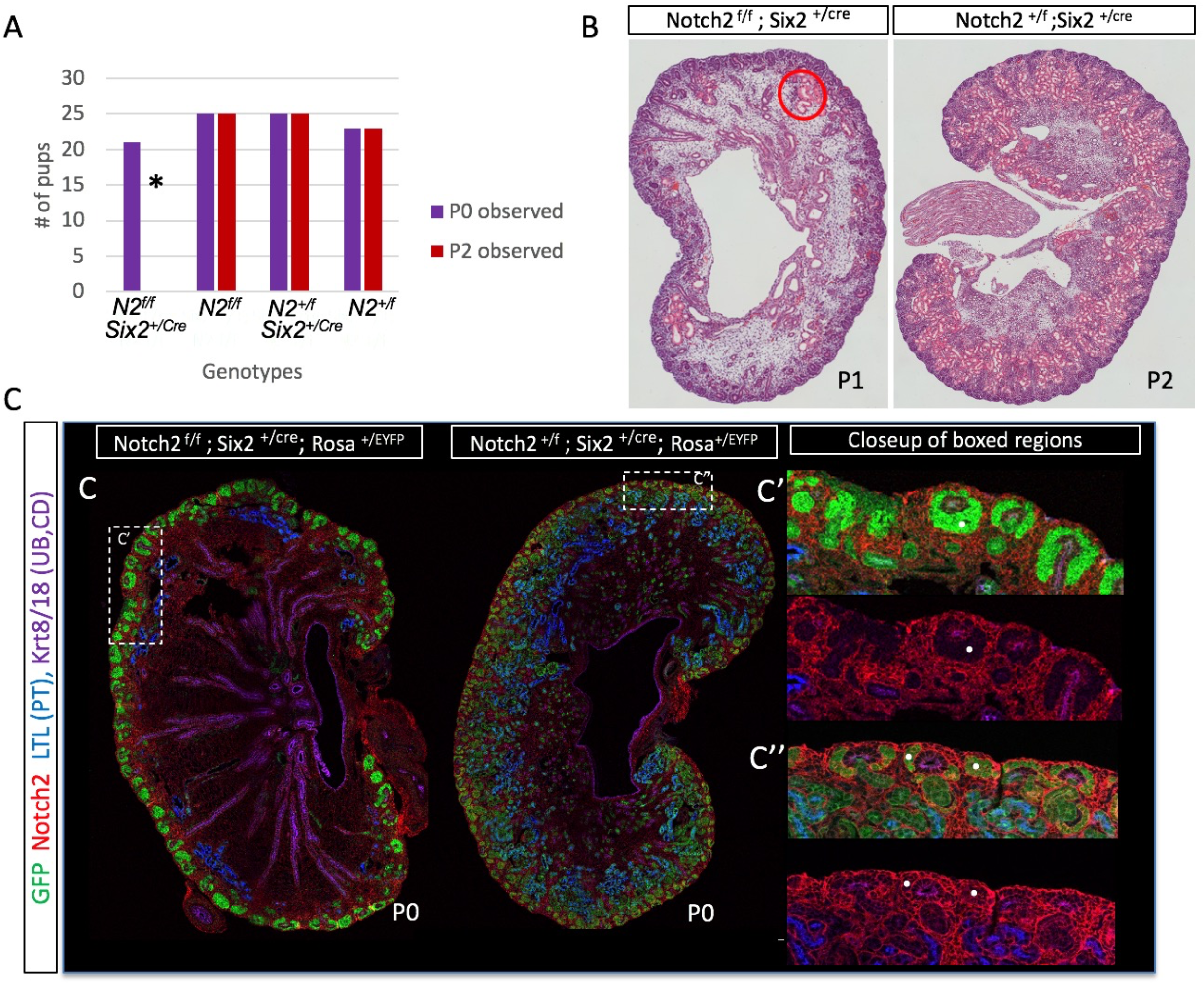
Survival, Histology, and immunostaining of mice with Notch2 deficient nephrons. **A**: Survival analysis. Six2^+/Cre^ Notch2^f/f^ mice were born at normal mendelian ratios, but none survived to P2. **B**: H&E stains of Six2 mutant and wild type kidneys from P1 and P2 respectively. The red circle indicates an area of PT cyst formation in the mutant. **C:** Six2 mutant and wild type kidneys from litter mates euthanized at P0 were stained with GFP (lineage trace), Notch2, LTL (PT marker) and Krt8/18 (UB and CD marker). Dashed white boxes were enlarged on C’, C”. **C’**: (top) View of the marginal zone region from a Six2^+/Cre^ Notch2^f/f^ with GFP (green) activated by Cre, and Notch2 (red) detected only in surrounding, GFP-negative cells. (bottom) Same image as above, showing only the Notch2 stain. White dot identifies similar structures in both panels. **C”**: (Top) View of the marginal zone region from Six2^+/Cre^ Notch2^+/f^ (wild type) with GFP (green) activated by Cre, and Notch2 (red). (Bottom) Same image as above, showing only Notch2. White dot identifies similar structures in both panels.

Hematoxylin-eosin (H&E) staining of postnatal day 1 (P1) *Six2*^+*/Cre*^*; Notch2^f/f^* and P2 *Six2*^+*/Cre*^; *Notch2*^+*/f*^ control littermates showed greatly diminished nephrogenesis in the mutants (Figure 2B, note the few nephrons forming due to mosaicism) indistinguishable from the description in (Cheng et al., 2007). Immunofluorescence performed on P0 kidneys from *Six2*^+*/Cre*^*; Notch2^f/f^*; *Rosa*^+*/EYFP*^ and *Six2*^+*/Cre*^*; Notch2*^+*/f*^; *Rosa*^+*/EYFP*^ controls detected GFP (lineage trace), Notch2, LTL (PT marker) and Krt 8/18 (UB and collecting duct (CD) marker) with the specified antibodies (Methods). As reported earlier, some proximal tubules were identified in *Notch2*-deficient kidneys, but only a few cells were marked with GFP, suggesting that these epithelial cells arose from *Notch2* expressing NPC. By contrast, nearly all nephron epithelia and NPC in control mice were positive for GFP (Figure 2C). Notch2, abundant in controls, (Figure 2C”) was not detected in NPC of GFP+ epithelia of *Six2*^+*/Cre*^*; Notch2^f/f^*; *Rosa*^+*/EYFP*^ kidneys (Figure 2C’). Finally, to ask if the milder phenotype reported previously reflected inefficient Cre-mediated recombination, we generated *Fgf20*^+*/Cre*^*; Notch2^f/f^* pups. Fgf20 is expressed at low levels in NPC and other epithelia such as the skin appendages and mammary glands (Barak et al., 2012; Elo et al., 2017; Huh et al., 2020; Huh et al., 2013; Huh et al., 2015; Yang et al., 2018), and our previous analyses suggested it is less efficient than Six2TGC. *Fgf20*^+*/Cre*^*; Notch2^f/f^* mice were born at normal Mendelian ratios, survived with no physical impairments, and were used for breeding. H&E staining of *Fgf20*^+*/Cre*^*; Notch2^f/f^* mice at P0 identified cysts in the kidneys with significantly fewer nephrons than controls (Figure 3A-B). The survival of *Fgf20^+/Cre^; Notch2^f/f^* mice, the low nephron numbers, and the appearance of cysts were all reminiscent of what we reported for the *Six2*^+*/Cre;*^ *Notch2^f/f^* mice at WUSM (Surendran et al., 2010a; Surendran et al., 2010b). This is consistent with the idea that the deletion efficiency of Cre in Six2TGC mice is sensitive to unidentified environmental factors.

**Figure 3:**
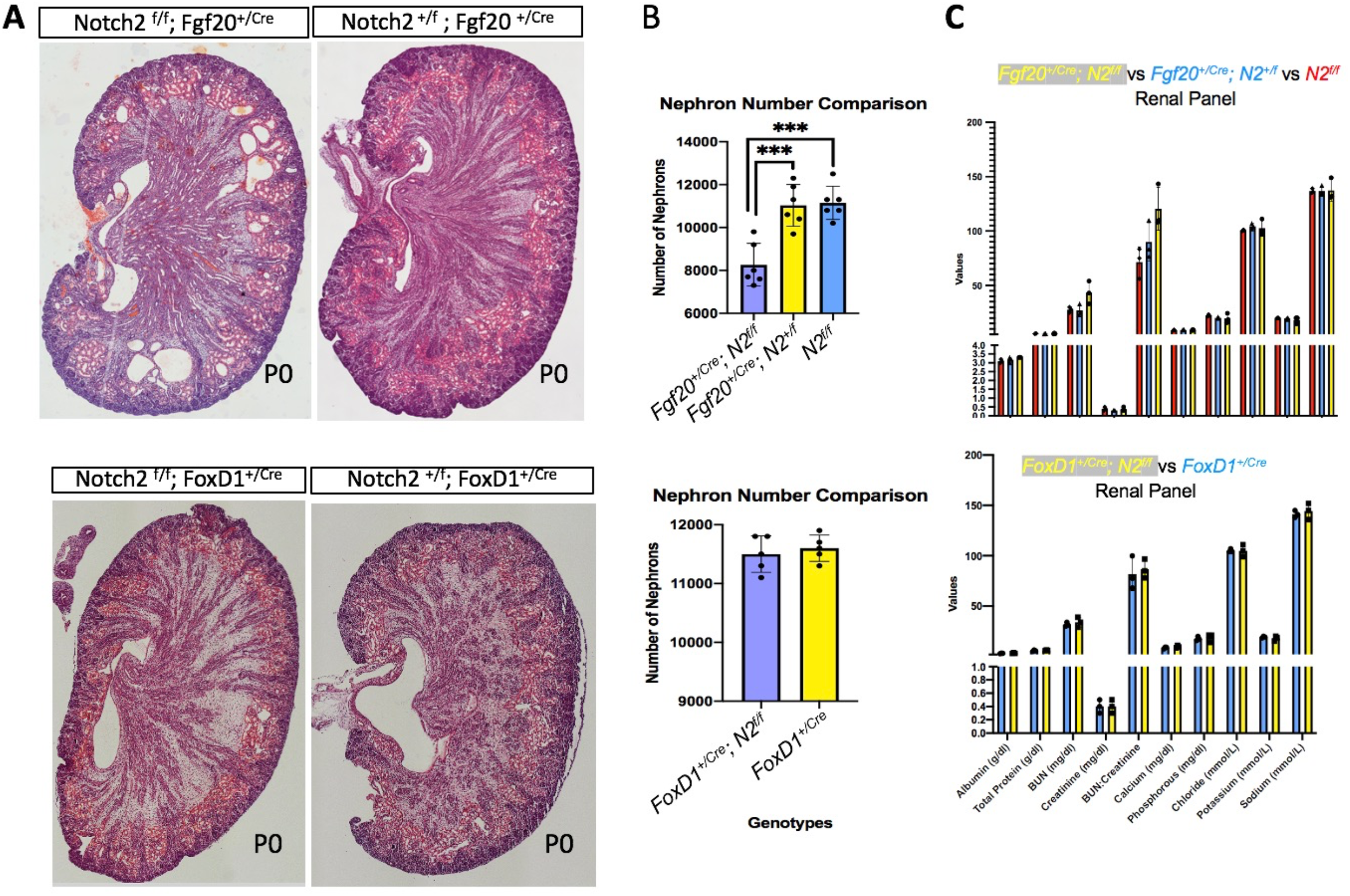
Histology, nephron counts and renal function analysis in Fgf20^+/Cre^ and FoxD1^+/Cre^ Notch2-deleted kidneys. **A.** Histological analysis of *Notch2 ^f/f^; Fgf20*^+*/Cre*^ and *Notch2 ^f/f^; FoxD1*^+*/Cre*^ kidneys. **B.** Nephron counts in these strains. Data is presented as sample group mean ± s.d. (n = 6 kidneys per group for Fgf20^+/Cre^;N2^f/f^, Fgf20^+/Cre^;N2^+//f^, and N2^f/f^; n = 5 kidneys per group for FoxD1^+/Cre^;N2^f/f^ and FoxD1^+/Cre^). For the Fgf20 group, statistical significance was evaluated by one way ANOVA with Tukey’s multiple comparison tests, *** denotes p ≤0.001. For the FoxD1 group, statistical significance was evaluated by unpaired two-tailed t-test, not significant, performed in GraphPad Prism 8. and **C.** Renal function in these strains after P90; n=3 for *Fgf20*^+*/Cre*^*;N2^f/f^*, *Fgf20*^+*/Cre*^*;N2^f/f^*, *FoxD1*^+*/Cre*^*;N2^f/f^ FoxD1*^+*/Cre*^, and *Fgf20*^+*/Cre*^*;N2^f/f^*. S.D represent standard deviation of the mean. All Multiple tTest and raw data are presented in Supplemental Table S1. All Multiple tTest adjusted values were >0.5.

To ask if loss of *Notch2* in the stroma or reduced nephron numbers had untoward effect on long term renal health, we tested renal function in *FoxD1*^+*/Cre*^*; Notch2^f/f^* and *Fgf20*^+*/Cre*^*; Notch2^f/f^* mice aged between 60 and 90 days. *FoxD1* is expressed in the stroma, and later in podocytes (POD) (Boyle et al., 2014). Like *Fgf20*^+*/Cre*^*; Notch2^f/f^* mice, *FoxD1*^+*/Cre*^*; Notch2^f/f^* mice were born at normal Mendelian ratios, survived with no physical impairments, and were used for breeding. *FoxD1*^+*/Cre*^*; Notch2^f/f^* kidneys and nephron numbers appeared indistinguishable from controls (Figure 3B), as reported earlier (Boyle et al., 2014). To measure renal function, 0.5 ml of blood were collected from three individuals per genotype (*FoxD1*^+*/Cre*^*; Notch2^f/f^*; *FoxD1*^+*/Cre*^*; Notch2*^+*/f*^ and *Fgf20*^+*/Cre*^*; Notch2^f/f^*; *Fgf20*^+*/Cre*^*; Notch2*^+*/f*^*; Notch2^f/f^*) and sent to a diagnostic laboratory (IDEXX) for analysis. No significant differences between mutant and wild type litter mates were detected, indicating normal renal function in these strains (Figure 3C, Supplemental Table S1). Thus, the threshold for formation of a kidney with sufficient function to support viability can be satisfied with a delay in *Notch2* deletion timing.

### Quantifying Fate Choices of Notch2 Deficient Cells Confirms Preferential Loss of Proximal Cell Fates

To characterize and quantify the cell types generated by *Notch2*-deficient NPCs, mutant and control kidneys were stained with antibodies to Tfap2b, an early distal tubule (eDT) marker, GFP, a lineage tracer, and the lectin LTL, a proximal tubule (PT) marker. Independently, we stained adjacent sections with antibodies to GFP, LTL, and Hnf4a (an early proximal tubule marker; ePT) to confirm that we did not miss any Hnf4a+ populations in Notch2 deficient kidneys (Supplementary Figure S2). The tissue was imaged on a confocal microscope, and captured in Imaris (Figure 4A). A control littermate stained with the same antibodies demonstrates that all nephron epithelia were derived from NPC in which GFP was activated following Cre-mediated recombination (note the transition from LTL to Tfap2b in the LOH, Figure 4B). The stained *Notch2*-deficient tissue from three individuals was initially divided into ~20 arbitrary sectors radiating from the papilla, totaling 71 sectors in three mutant kidneys (Figure 4A, Supplemntal Table S2). Three observers (one blinded to the experimental goal) counted the sectors that contained only LTL, LTL&GFP, only Tfap2b, Tfap2b&GFP, or LTL&Tfap2b, counting the latter even if just one LTL-positive cell was detected. The counts were averaged to determine standard deviation. The percentage of sectors that contained Tfap2b+ DT (78.4%) was significantly higher than the percentage that contained LTL+ PT (58.69%; Figure 4C, *p*=0.013). Sectors containing Tfap2b+ and LTL+ cells were considered GFP positive if 50% or more of the cells in the cluster were GFP+. By that definition, a significantly higher percentage of DT sectors were GFP+ than PT sectors, 44.6% to 13.6% respectively (Figure 4C, *p*=0.007), suggesting that in the absence of Notch surviving epithelia are distal in character.

**Figure 4:**
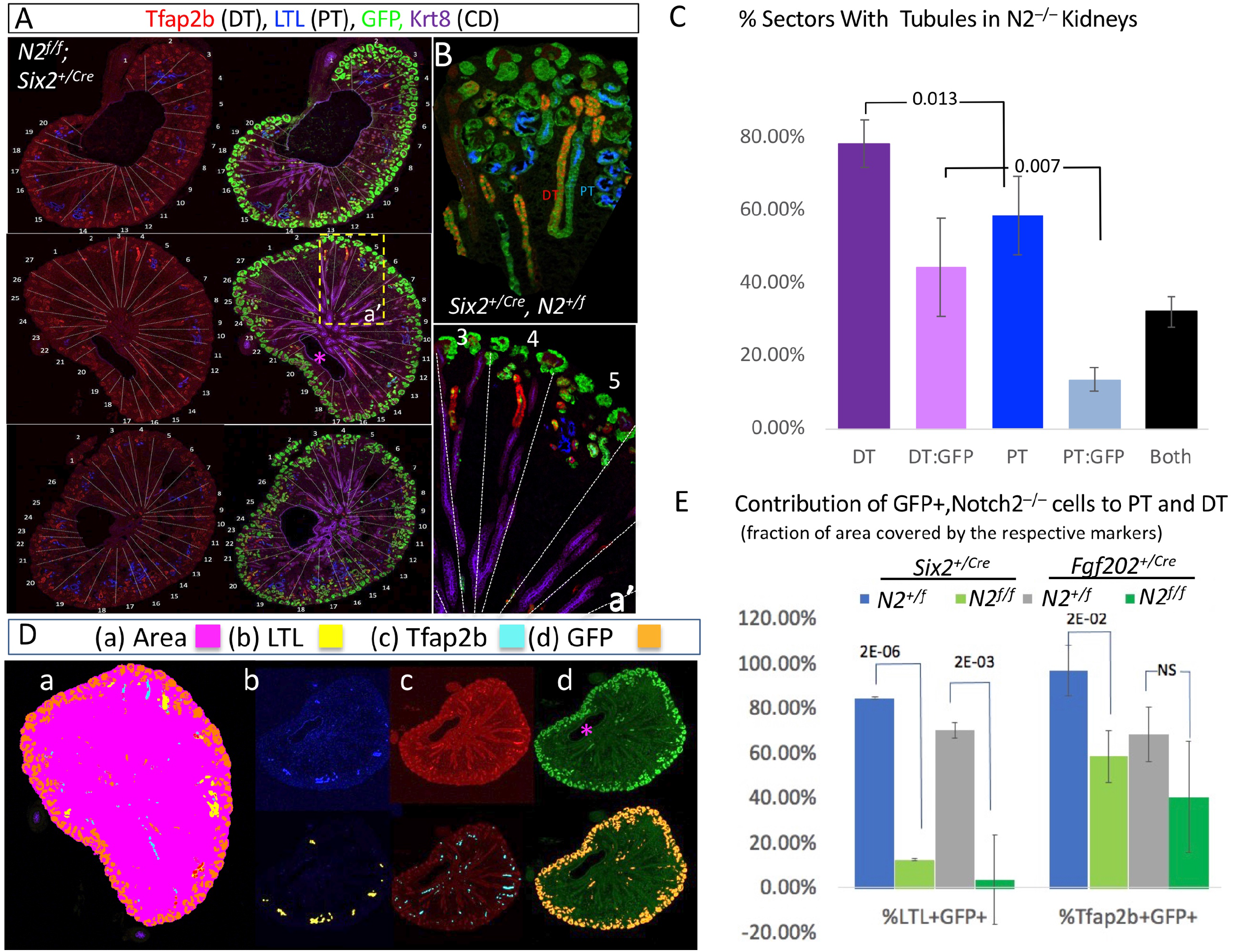
Notch signals are necessary to promote PT fate. **A.** Kidneys stained for GFP (linage tracer), LTL (PT) and Tfap2b (DT) were divided into 20 or 21 arbitrary sectors in kidneys from three animals (n=71 sectors; see Supplemental Table S2). A boxed region from one kidney is shown in a’. **B.** Close up of wild type kidney. **C.** The 71 sectors shown in (**A**) were scored by three independent observers for the presence or absence of any of the three markers and averaged. **D-E.** To obtain quantitative data the kidneys were imaged in IMARIS, segmented by color (**Db-d,** top), and the area deemed positive for each color presented in psudocolor. In **Da,** the total kidney area (pink, asterisk in **Dd** marks the section used for this determination) and all the color combination are presented. The area positive for each fate marker was calculated by the software as a fraction of the total kidney area and plotted in **E**.

To provide a more accurate accounting of fate choices we created a binary algorithm in Nikon Elements to measure the kidney area (Figure 4Da), the area covered by PT (LTL+, Figure 4Db), the area covered by DT (Tfap2b+, Figure 4Dc), and the area covered by GFP (Figure 4Dd). Using these masks, we calculated the area that was LTL+/GFP+ and Tfap2b+/GFP+ in μm^2^, and extracted its percentage from the total DT and PT areas (Figure 4E). In control, *Notch2*^+*/f*^ littermate kidneys, 86.6% of LTL+ and 96.8% of Tfap2b+ cells were also GFP+ (Figure 4E, Supplemental Table S2). In *Notch2*-deficient kidneys, only 12.4% of LTL+ cells were GFP positive, in contrast with 58.5% of Tfap2b+ cells that were also GFP positive. Thus, *Notch2* deficient cells are more than four times as likely to express the early DT marker than they are to express the PT marker. A similar choice bias was observed in *Fgf20*^+*/Cre*^; *Notch2^f/f^* kidneys, with only 3.4% of LTL+ cells being GFP+ against 40.3% of Tfap2b+ expressing GFP. *Fgf20*^+*/Cre*^; *Notch2^f/f^* kidneys have significantly more GFP-LTL+ cells than the *Six2*^+*/Cre*^; *Notch2^f/f^* kidneys (Figure 4E), suggesting that a later deletion may allow PT cells to survive w/o Notch. It should be noted that although *Six2*^+*/Cre*^; *RBP^f/f^* kidneys contain very few epithelial cells, 42.9% of 71 sectors were *Tfap2b*+; *LTL*− but only 0.42% that was *LTL* +; *Tfap2b* – (*p*=0.014; Supplementary Figure 3A-B, Supplementary Table S2). Notably, and in contrast to *Wnt4*^+*/Cre*^; *Notch1^f/f^*; *Notch2^f/f^* (Chung et al., 2017), *Wnt4*^+*/Cre*^; *RBP^f/f^* kidneys were indistinguishable from *Wnt4*^+*/Cre*^; *RBP*^+*/f*^ kidneys (Supplemental Figure 3C), suggesting that perdurance of RBP protein (Turkoz et al., 2016) may suffice to complete nephrogenesis. *Wnt4*^+*/Cre*^; *RBP^f/f^* mice did not survive (Supplemental Figure 3D), however, perhaps due to the requirement of Notch signaling in other Wnt4 expressing tissues (Caprioli et al., 2015), a possibility we did not pursue further. These analyses complement our earlier published conclusion, inferred based on the patterns of LTL and Cdh1 (Cheng et al., 2007), that Notch2 expression guides cells towards a proximal fate before acting in all cells to promote proliferation/maturation.

### The Stabilized Notch1 Intracellular Domain expands PT at the expense of NPC and DT

When N1ICD^ΔPEST^ was over expressed in Six2-expressing progenitors, it drove epithelialization in the absence of Wnt4 and Wnt9b, producing mostly LTL+, Cdh1− and Wt1+ cells (Boyle et al 2011, Cheng et al 2007). To ask if early DT cells are also present under these conditions we crossed a *Six2*^+*/Cre*^ sire with *Rosa^NICD/NICD^* dams (Murtaugh et al., 2003) and analyzed nephrogenesis histologically at E15.5, as we did previously (Boyle et al 2011, Cheng et al 2007), using additional markers and obtaining bulk RNA-seq data. The kidneys were isolated from the embryos and processed for whole mount immunofluorescence with antibodies against Wt1 (NPC and podocyte marker), Hnf4a (PT marker), Tfap2b (DT marker) and GFP (co-expressed with N1ICD^ΔPEST^). We again noted large expansion of epithelia and paucity of Six2+ NPC but could only detect a few cells that expressed Tfap2b (Supplementary Movies S1, S2). Note they were also positive for Wt1, LTL, and Hnf4a (Supplementary Movie S1). GFP+/LTL+/Hnf4a+ triple positive cells dominated (Figure 5A-C, Supplemental Movies S1, S2). Interestingly, Wt1+ positive cells did not stain for GFP, suggesting they may have lost NICD expression, or arose from cells in which NICD was not activated by the Six2TGC allele. The lineage tracing by Chung et al, 2017 supports the former hypothesis.

**Figure 5:**
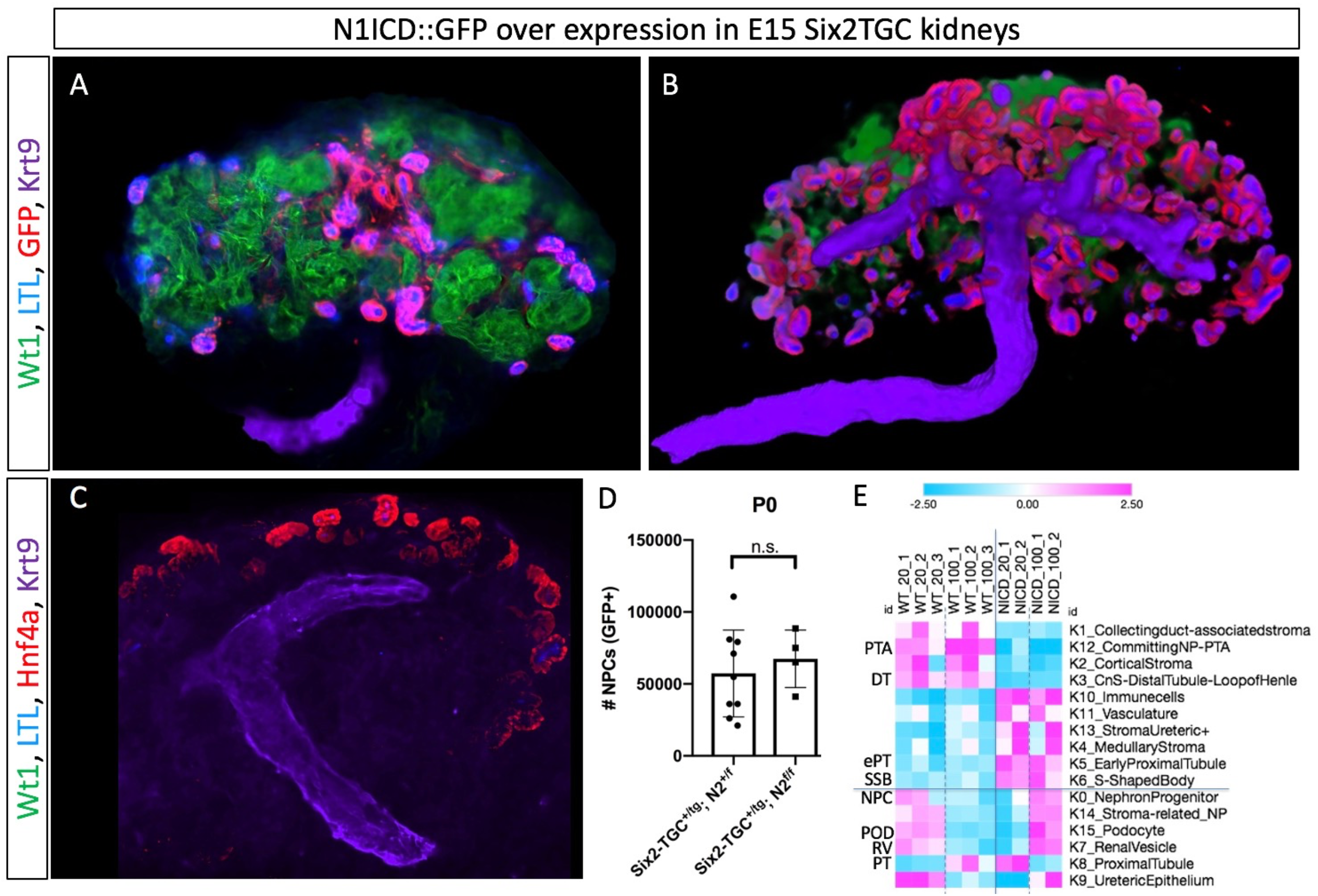
N1ICD^ΔPEST^ compels differentiation, proximal development at the expanse of distal cells. **A.** Whole mount of a kidney anlage expressing N1ICD^ΔPEST^. Note the Wt1+ cells (green) are not expressing GFP (red). **B.** A frame from Supplemental Movie S1. **C.** A frame showing Hnf4a expression from Supplemental Movie S2. **D.** P0 kidneys with the genotypes noted were dissociated and GFP+ cells (Six2+ NPC) counted by flow cytometry. Data is presented as sample group mean ± s.d. (n = 9 control Six2-TGC^+/tg^;N2^+/f^ and n = 4 experimental Six2-TGC^+/tg^;N2^f/f^ isolated from two litters), n.s. denotes non-significant p-value (p=0.5561) by unpaired two-tailed t-test performed in GraphPad Prism 8. **E.** Bisque deconvolution assigned relative cell type enrichment in bulk RNA data samples from three wild type and two N1ICD^ΔPEST^ expressing kidneys using the top 20 or 100 markers from scRNA dataset (Combes et al., 2019) as a standard.

The rationale for this study stems in part from past practices of inferring fate outcomes from a limited number of “marker” genes. To consider the broadest possible number of transcripts we analyzed differential gene expression induced by stabilized N1ICD^ΔPEST^ in two N1ICD^ΔPEST^over-expressing kidneys and three control littermates. mRNA was isolated from each kidney anlage and submitted to Novogene for amplification, library construction and sequencing. 5,283 transcripts were differentially expressed (Supplementary Table S3). Among the 10% most underrepresented genes in the N1ICD^ΔPEST^ over-expressing kidney were *Calb1* (LogFC −9; UB), *Six2* (−8.7; NPC), *Slc12a1* and *Slc12a3* (−8.26 and −8.16, respectively, DT/LOH), *Sal3* (−6.6, NPC, DT/LOH), and *Fgf8* (−6, NPC, DT/LOH). *Tfap2b* (−5, DT/LOH) and *Sim1* (−4.7, DT) were in the next decile, ranking 56th and 65th respectively. *Hp* (7.4, ePT) was present among the most over-represented decile, with Notch targets *Cdh6* (1.6) and Nrarp (0.58) mildly over expressed (Supplementary Figures S4, S5 and Supplementary Table S3). The near elimination of *Six2* and other NPC genes (Supplementary Figure S5A) was consistent with the proposal that NICD repressed Six2 (Fujimura et al., 2010). To test if the absence of Notch delayed niche exit and thus led to expansion of NPC, we sorted GFP+ cells from kidneys isolated from four P0 *Six2*^+*/Cre*^; *Notch2^f/f^* and nine *Six2*^+*/Cre*^; *Notch*^+*/f*^ pups. No significant difference was observed in the number of GFP+ NPC (Figure 5D), inconsistent with the proposed role for Notch as a Six2 repressor regulating niche exit. Notably, while Notch activation clearly can drive *Six2*-expressing cells to epithelialize, none of the known Notch targets were found in Pretubular Aggregate (PTA) in our (Jarmas et al., 2021) or in published (Combes et al., 2019) datasets, suggesting that it functions after the epithelia form, perhaps promoting proliferation (Supplementary Figure S6) and maturation.

Next, we manually annotated 2683 transcripts based on a published scRNA-seq dataset (Combes et al., 2019) to plot the changes in gene expression as a heatmap segregated by assigned cell type (Supplementary Figure S5A and Supplementary Table S3). NPC and DT/LOH genes were underrepresented in N1ICD^ΔPEST^ relative to control, while ePT, POD and inflammation markers were overrepresented in N1ICD^ΔPEST^ expressing kidneys. Notably, mature PT transcripts declined in N1ICD^ΔPEST^ overexpressing cells. To gain a more quantitative insight, we performed deconvolution analysis of the bulk data with BisqueMarker (Jew et al., 2020) applying a weighted PCA approach using either the top 20 or the top 100 markers from (Combes et al., 2019) (Figure 5E, Supplementary Table S4). Based on this analysis, PTA and DT were under-represented, while SSB and ePT were overrepresented in N1ICD^ΔPEST^ overexpressing kidneys. NPC, POD, RV, and mature PT were underrepresented in the top 20 marker deconvolution and overrepresented using the top 100 markers, perhaps reflecting an abnormal redistribution of transcripts in these cells consistent with the supraphysiological presence of N1ICD^ΔPEST^. In aggregate, results from manual curation and regression-based computational analyses are most consistent with a role for Notch in supporting selection of the proximal fates early, before promoting epithelial proliferation and maturation of both distal and proximal tubule cells.

### NOTCH Signaling Required for Proximal Fates to Emerge in Human Kidney Organoid

To test the idea that a Notch function in nephron segmentation is conserved among mammals, we targeted the second exon of the *Nicastrin* (*NCSTN*) gene in the NHSK hiPSC line with gRNA and CRISPR-Cas9 (Figure 6). *Nicastrin* is a non-redundant member of the γ-secretase complex, and in its absence, this enzyme is highly unstable, resulting in compete loss of signaling from all four Notch receptors (Li et al., 2003; Nguyen et al., 2006). GFP (and thus, CRISPR and gRNA) expressing cells were sorted, and colonies screened for NCSTN expression and Notch activation (Figure 6B-C). DNA from two *NCSTN* deficient clonal cell lines was sequenced and found to contain deletions resulting in a frame shift and termination in Exon 2 (Clone 1) or 3 (Figure 6A). Kidney organoids were generated from Clone 20 (C20) as control that underwent CRISPR transfection without losing *NCSTN*, and C1 *NCSTN*-/- cells using a combined Morizane and Takasato protocols (see methods, (Morizane and Bonventre, 2017; Morizane et al., 2015)). Organoids were analyzed for the expression of ePT, eDT and UB markers by confocal imaging and Z stacks created for each. In contrast to robust nephrogenesis induced in the C20 organoid by day 22, C1 *NCSTN*-/- organoids were smaller, and contained few clusters of KRT8/18 positive cells connected to TFAP2b-expressing tubules. PODXL, HNF4A or LTL expressing cells, abundant in the parental line, were not detected (n=4 separate differentiation experiments, Figure 6, Supplementary Movie 3). This observation indicates that Notch function in P/D axis segmentation is conserved across mammalian species.

**Figure 6:**
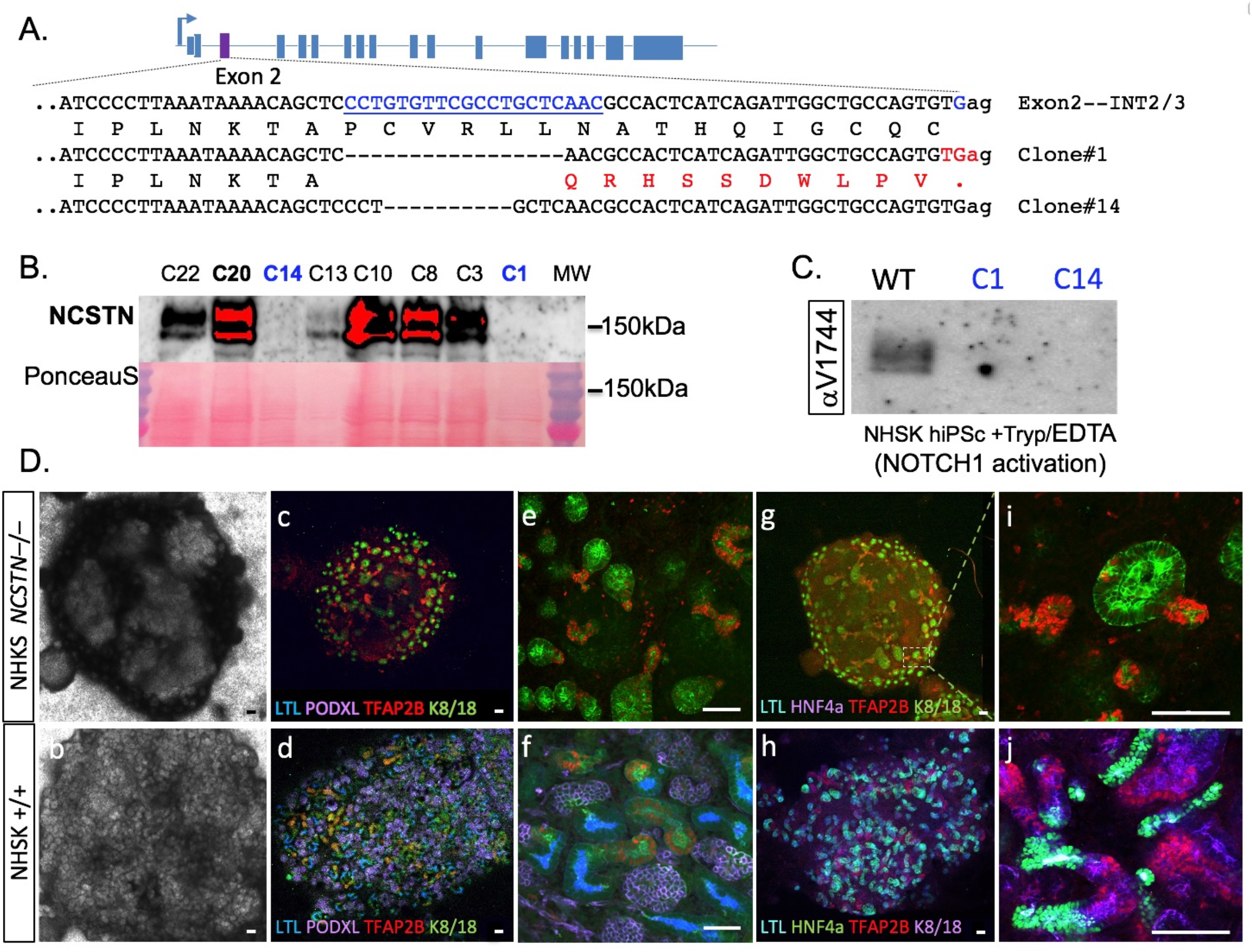
CRISPR-CAS9 mediated NCSTN mutant iPSCs generate kidney organoids composed solely of immature distal tubular epithelia. **A.** Schematic representation of NCSTN gene disruption by CRISPR/Cas9. The deleted nuclotides in exon 2 are marked with blue and the resulting frameshifted amino acids and premature stop codon in the two mutated clones (#1 and #14) are marked in red. **B.** Western blot analysis for NCSTN expression in unmodified clones (C22, C20) and NCSTN-/- clones (C1, C14) showing loss of NCSNT protein expression in the latter. **C.** NOTCH1 activation with Tryp/EDTA in an unmodified control (C20) and the two NCSTN-/- clones (C1 and C14) was anlyzed by Western blot probed with the αVal1744 rabbit monoclonal antibody, detectng the VLLS epitope of N1ICD. N1ICD was detected in C20 but not in either of the NCSTN-/- clones, indicting loss of γ-secretase. **D.** hiPSC-derived kidney organoids from C20 (unmodified control; lower panel) and C1 (NCSTN- /-; upper panel) show significantly lower percentages of differentiated structures in the NCSTN-/- compared to controls in bright field images of kidney organoids derived from NCSTN-/- **(a)** and unmodified control **(b)** iPSC clones. c-h; Whole mount immunofluorescent staining for early proximal nephron structures (i.e. HNF4a, LTL and PODXL) and early distal nephron structures (i.e. TFAP2B, KRT8/18) demonstrating expression of distal nephron markers but not expression of proximal nephron markers in the NCSTN-/- **(c, e, g)** in contrast to unmodified control **(d, f, h)**. Both bright field and IF images are representative of 4 separate kidney organoid differentiation experiments for each cell clone. 8-10 organoids from every clone in each differentiation experiment were immunostained for these markers. Scale bar=100μm.

## Discussion

Investigations into the developmental role of Notch signaling in the mammalian kidney began at the turn of the 21st century, with identification of *Jag1* mutation in Alagille syndrome (Heritage et al., 2000) and concurrent studies in model organisms (McCright et al., 2001). The importance of this pathway to the completion of nephrogenesis is broadly accepted, but whether Notch signaling had a role in nephron segmentation is disputed. Here we establish that indeed, prior to its roles in maturation and proliferation, Notch signals contribute to form the proximo-distal axis in the emerging nephron epithelia.

Using Notch pathway loss of function models investigators observed that MET can occur, albeit at reduced rates, and noted paucity of epithelial cells but failed to provide a definitive mechanistic explanation for this shortage. Notch promotes epithelial proliferation in a Jag1 dependent process (Cheng et al., 2007; Ungricht et al., 2022) and perhaps, survival (one report noted increased apoptosis in the marginal zone mesenchyme of *Presenilin1* hypomorphs at E15.5 (Wang et al., 2003)). Interestingly, Jag1 both promotes proliferation in trans (on neighboring cells) and inhibits it in cis (where it is expressed), the latter function only uncovered by screening chimeric organoids (Ungricht et al., 2022). There is also agreement that Notch signals most likely stabilize and/or promote differentiation of all epithelial nephron fates, initially proposed by (Fujimura et al., 2010) and elaborated in (Chung et al., 2016; Chung et al., 2017). Notch is not likely to contribute to selection of differentiation vs. self-renewal in the NPC: if Notch arbitrated this choice via lateral inhibition (Bray, 1998), without Notch cells would be expected to assume an alternative fate. Accordingly, it was suggested that Notch inhibits Six2 (Chung et al., 2016; Fujimura et al., 2010), and that Six2-expressing NPC accumulated in Notch-deficient kidneys. However, the same number of NPC could occupy a larger area in the smaller Notch-deficient kidneys and indeed, we show that the loss of Notch signaling did not lead a change in the number of *Six2*+ progenitors (Figure 5D). Whereas Notch is not acting to switch cells from self-renewal to differentiation, Notch signals could promote proximal development within a developmental window, defined pharmacologically with γ-secretase inhibitors (Cheng et al., 2003; Cheng et al., 2007). This role would predict that if Notch contributed to segmentation, in its absence, a skewed fate map will be observed. Testing this prediction produced results that varied based on the choice of markers used and can either support a role in segmentation (Cheng et al., 2007), or refute it (Chung et al., 2016; Chung et al., 2017). In addition, the broad domain in which Notch2 was deleted by the Pax3-Cre used in the Cheng study, the inability of N2ICD or of late expression of N1ICD^ΔPEST^ to promote proximal fate (Chung et al., 2016; Chung et al., 2017; Fujimura et al., 2010), and the absence of mature PT or DT markers after deleting Notch1/2 in RV epithelia (Chung et al., 2017) further diminished confidence in the proposed role for Notch in segmentation.

The emergence of single cell transcriptomics revealed that earlier studies addressing segmentation were deficient in selecting the markers for their analysis. Whereas Cheng and colleagues argued that broad epithelial marker *Cdh1* can be used to identify distal cells, they did not buttress that claim with any additional distal markers. The counterargument relied on the absence of other markers (Chung et al., 2017) which we now realize reflect mature distal and proximal fate. A pan-epithelial block in maturation prevents expression of these markers, and their absence does not exclude a Notch function in segmentation operating before maturation.

Our analysis of the single cell transcriptome of developing kidneys revealed a hierarchy of distal and proximal markers enabling a more careful, higher resolution reexamination of Notch functions during nephrogenesis. Lineage analysis in two loss of function mouse models, the Six2-Cre and Fgf20-Cre, or in NICASTRIN-deleted human iPSC-derived kidney organoid model, reveals that these cells give rise to predominantly Tfap2b+ cells, a transcription factor committing cells to the distal/LOH fate, connected to KRT8/18 positive connecting duct cells (Marneros, 2020; Naganuma et al., 2020). We demonstrated that lower levels or delayed Cre expression in mouse kidneys is compatible with survival since enough nephrons form, suggesting that at WUSM, Six2-Cre activity resembled Fgf20-Cre in timing and/or amount. Importantly, although urine analysis did not reveal any renal dysfunction, the proximal/distal ratio was altered in *Fgf20-Cre*, *Notch2^f/f^* kidneys. Based on this model, we anticipate that analysis of archival kidneys from Alagille patients will show a similar bias with fewer PT cells relative to DT/LOH.

Correspondingly, marker and transcriptome analyses of arrested anlage expressing a stabilized N1ICD^ΔPEST^ demonstrated a skewing towards immature early epithelia with elimination of NPC and great reduction in DT-fated cells. Notably, despite the strong bias towards the early proximal fate, *Lhx1*, thought to be a mediator of Notch activity (Chung et al., 2017), was among the transcripts suppressed by this hyper-physiological Notch signal, as was the Notch target gene *Hes5* (supplemental Figure 4A); Hnf1ß levels were unchanged. The abundant Wt1+ cells, derived from Cre deletion (i.e., marked by the lineage tracer, as in (Chung et al., 2017)), were negative for GFP co-expressed with N1ICD^ΔPEST^, consistent with *in vitro* observations (Boyle et al., 2011) and with the hypothesis that Notch hyperactivation is inconsistent with podocyte development (Cheng et al., 2007). Finally, and in contrast to the outcome when *Notch2* (herein) or *Notch2 and Notch1* genes are deleted with Wnt4-Cre (Chung et al., 2017), deletion of *RBP* using the Wnt4 Cre driver had no impact on nephron development, suggesting that by the time the long-lived RBP protein (Turkoz et al., 2016), has been depleted, all Notch functions in the NPC lineage have been fulfilled.

How can the various studies reviewed herein be reconciled into a single unifying model for Notch function? We arranged the published results along a developmental ruler (Figure 7). Analyses of Notch2 loss of function induced with either Pax3-Cre or Six2-Cre show that Notch2 was not required for proliferation of naïve or committed NPC, nor was it required for *Wnt4* activation (Boyle et al., 2011; Wang et al., 2003). The cells expressing *Notch2* and *Cited1* are unlikely to encounter the very few ligand expressing NPC within the same domain (Figure 1), and when they do, it may accelerate exit since we do not see marked NPC clones with the fate marker Notch2-Cre (Liu et al., 2013). These observations in mice are supplemented by Network analysis in human first trimester kidneys (Supplementary Figure 5B, (Lindstrom et al., 2021)), consistent with Notch activity emerging after MET, in the PTA, where Notch signaling shifts into gear and promotes early segmentation into ePT (it is not required for the eDT fate). In all nephron epithelia Notch signals are required for maturation, proliferation and/or survival (Chung et al., 2016; Chung et al., 2017; Fujimura et al., 2010; Ungricht et al., 2022) prior to or during the formation of the S-shape body (*RBP* deletion in this study, and pharmacological analysis (Cheng et al, 2003)). One consequence of the maturation-promoting Notch function is that cell fate choices in loss-of-function models cannot be analyzed with late, differentiation-dependent lineage markers (whose expression is lost); this function can be detected by transcriptomic analysis or by use of early markers (presented in Figure 1, Supplementary Figure S1, and Naganuma et al., 2020). Deleting both receptors with Wnt4-Cre (acting in PTA/RV) is predicted to result in a similar bias, not detectable with the late markers used by (Chung et al, 2017). By contrast, using the same Wnt4-Cre line to remove both *RBP* alleles falls outside the Notch-sensitive window due to the perdurance of the RBP protein (Figure 7). When stabilized, hyper-physiological N1ICD^ΔPEST^ is over expressed from the weak Rosa26 locus in the NPC (using Six2-Cre or TAT-Cre), the pro-ePT, differentiation function of Notch is obvious, but maturation is impaired by sustained, strong Notch signals as evident by the reduction in mature PT markers. Since the NICD domain of Notch1 and Notch2 are interchangeable with no impact on kidney development (Liu et al., 2013), we favor the interpretation that lower amounts of N2ICD (Fujimura et al, 2010) are either insufficient to bias the outcome or do so in a manner requiring quantification of all transcripts.

**Figure 7:**
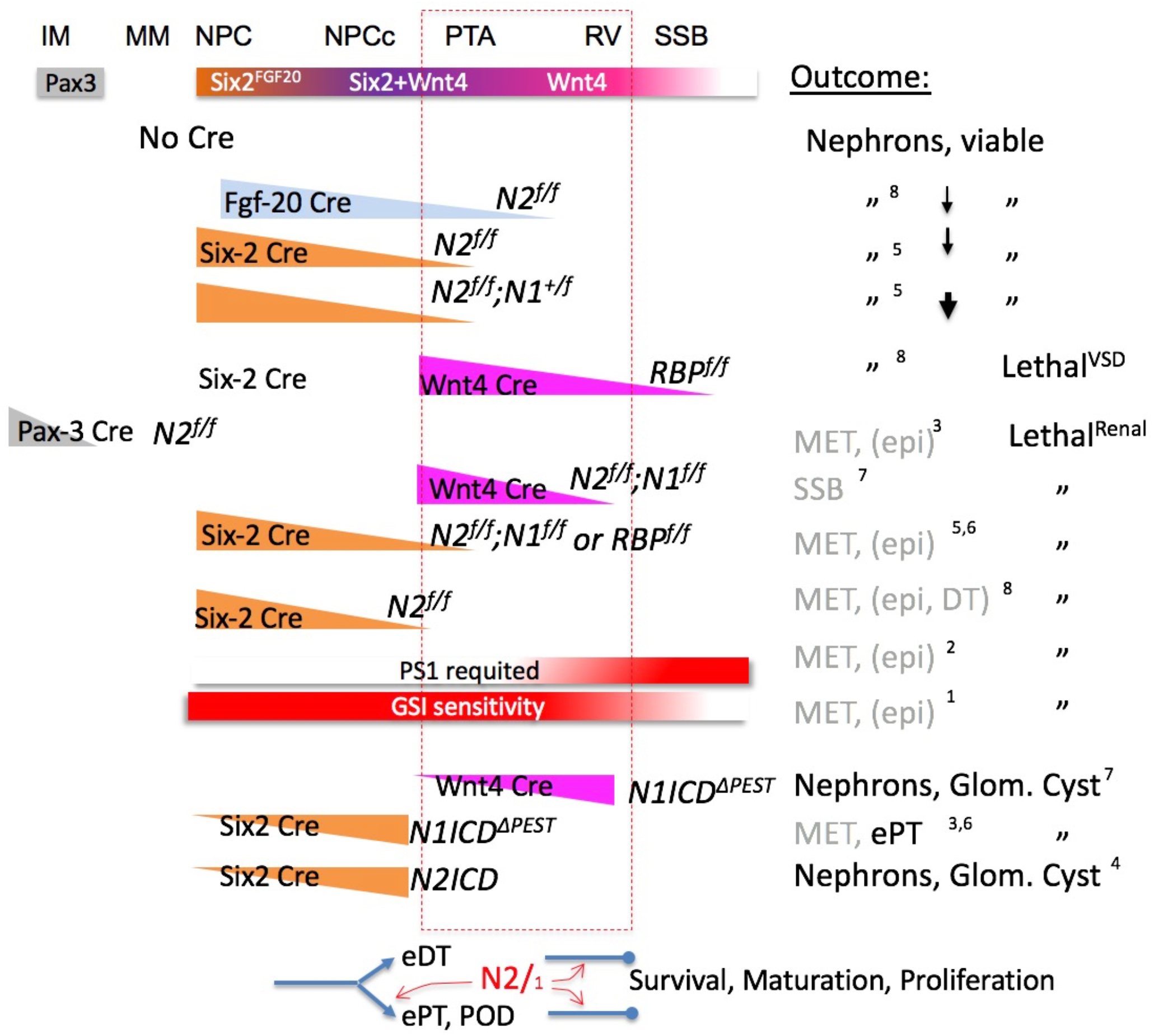
The temporal requirements for Notch signaling revealed genetic and pharmaceutical studies. Schematic representation of phenotypes along a developmental timeline (IM, intermediate mesoderm, MM, metanephric mesenchyme, NPC, nephron progenitors, NPCc, committed NPC; PTA, pretubular aggregate, RV, renal vesicle, SSB, S-shape body). Below the developmental timeline is a representation for the expected expression onset pf various Cre drivers (Pax3, Six2, FGF20, Wnt4). Of these drivers, FGF20 expression level is the lowest. The outcome column describes what is observed when conditional alleles, placed above the cre driver schematic, are removed. The triangular shape infers time to complete loss of protein function, graded boxes depict sensitivity to GSI (gamma-secretase inhibitor). The outcomes are presented in inverse order of severity (genetic loss of function first, then pharmacology, followed by over expression). Downwards pointing arrow indicates fewer nephrons than control (thin arrow= less significant than thick arrow). The temporal window is marked with a dashed red box. VSD; ventricular-sepal defect. Glom, glomerulus. Not included in the schema are the outcome in *NICASTRIN*-deleted organoids. The sources for the data in the figure: 1) Cheng et al, 2003; 2) Wang et al, 2003; 3) Cheng et al, 2007; 4) Fujimura et al, 2010; 5) Surendran et al, 2010a; 6) Chung et al, 2016; 7) Chung et al, 2017; 8) this study.

In conclusion, this study integrates two decades of Notch research in the metanephric kidney context to refine a timeline in which three Notch functions operate: 1) assistance in selection of ePT fate, 2) promotion of proliferation/survival of nascent nephron epithelia, and 3) their maturation, and extends these conclusions to the human. It is unclear if these Notch functions are separated by the S-phase, as they are during gonad development in *C. elegans* (Ambros, 1999), or by a cell division, as they are the *Drosophila* peripheral sense organs (Jan and Jan, 1994). The strength of Notch signals plays an important role, with weak/no early signal being compatible with the DT/LOH developmental trajectory but somewhat stronger signal needed for ePT, like the development of definitive hematopoietic stem cells in the AGM (Gama-Norton et al., 2015). All cells need an above-threshold signal to expand and mature. Too much (or persistent) signal, as in N1ICD^ΔPEST^, is deleterious (see also (Niranjan et al., 2008; Sweetwyne et al., 2015; Waters et al., 2008)). It remains to be determined which downstream effectors are involved in prompting the ePT fate, nephron epithelial expansion, and segment maturation.

## Materials and Methods

### Mice

mouse strains used were: *Foxd1^tm1(GFP/Cre)Amc^* (FoxD1-Cre) (Jax Stock No: 012463); *Six2^tm3(EGFP/Cre/ERT2)Amc^* (Six2TGC, *Six2*^+*/Cre*^; Jax Stock No: 009600); *Fgf20^tm2.1(Cre/EGFP)Dor^* (Fgf20-Cre)( Huh SH, et al. 2015); *Notch2^tm3grid^* (*Notch2*^+*/f*^) (Jax Stock No: 010525); *Rbpj^tm1Hon^* (*Rbpj*^+*/f*^) (Tanigaki et al., 2002), *Wnt4^tm3(EGFP/Cre)Amc^* (*Wnt4*^+*/Cre*^) (Jax Stock No: 032490) Rosa^NICD^ (Murtaugh et al., 2003), *Rosa^+/^* EYFP [Gt(ROSA)26Sortm1(EYFP)Cos],(Srinivas et al., 1999).

Mice were maintained at Cincinnati Children’s Hospital Medical Center animal facility following animal care guidelines. Our experimental protocols (IACUC 2018-0108/0107) were approved by the Animal Studies Committee of CCHMC. For embryonic experiments, embryonic day (E) 0.5 was noted at noon at the day the copulatory plug was observed. Genotyping was performed on toe or tail clips following standard genotyping protocols (Stratman et al., 2003). Oligonucleotides used are listed in Supplemtal Table S5.

### H&E staining

Kidneys were isolated from mice at P0, e15.5, e17.5 and e18.5. Once isolated, P0 kidneys were fixed in 4% PFA for 16-20 hours at 4°C and younger kidneys were fixed for 1 hour at room temperature. After washing 3X in PBS, kidneys were processed, embedded, and sectioned by CCHMC Pathology Core. Slides were then deparaffinized in xylene 3X for 2 minutes each and rehydrated in ethanol (100%, 70%, 50%, and 30%). Slides were then rinsed with distilled water. After staining with hematoxylin for 4-5 minutes, slides were rinsed in H2O, dipped once in 0.05% acetic acid, rinsed in H2O, dipped in 0.1% ammonium hydroxide 3X, rinsed in H20, and dipped in 80% ETOH 20X. Once stable in 80% ETOH they were dipped in eosin for 20 seconds, rinsed in an ethanol series to xylene before being mounted with mounting medium. Slides were left to dry o/n before being imaged on a Nikon NiE upright microscope.

### Immunofluorescent Staining

Kidneys were isolated and fixed as above. After washing 3X in PBS, kidneys were processed in an alcohol series xylene, embedded in paraffin, and sectioned by CCHMC Pathology Core. Slides were deparaffinized in xylene 2X for 10 minutes each, and rehydrated as above. Slides were then washed in PBS 3x for 5 minutes each, placed in a solution of Trilogy Antigen Retrieval (Cell Marque, 920P-10), and boiled in a rice cooker for 30 minutes. Slides were washed again in PBS 3x for 5 minutes each and then placed in a blocking solution [PBS, 5% Bovine Serum Albumin (BSA; Sigma-Aldrich, A8806-1G), 0.1% tween-20 (Fisher Scientific, BP337-500), and 10% normal donkey serum (NDS; Jackson Immu. Res. Lab, 017-000-121)] for 1 hour at room temperature. See Supplementary Table S5 for the list on antibodies used. Images were captured using a Nikon A1 inverted confocal microscope. Slides were incubated in secondary antibody for 1 hour at room temperature. Slides were then washed a final 3X in PBS-T for 30 minutes each, and mounted with Prolong Gold (Cell Signaling Technology, 9071S).

### Whole mount

Kidneys were removed from NICD mice at e15.5 and fixed in 4% PFA for 1 hour at room temperature, washed 2x with PBS and permeabilized and blocked for 3 hours with gentle rocking at room temperature in TSP-NS [PBS, 0.1% Triton 100X (Fisher Scientific, BP151-500), 0.05% saponin (Fluka, 47036), and 10% normal donkey serum]. Whole kidneys were submerged overnight in TSP-NS with specified primary antibodies (Supplemental Table S5) at 4°C with gentle rocking. Post incubation the kidneys were washed 3x for 10 minutes each in TSP at room temp. Secondary antibodies were diluted in TSP-NS (Supplementary Table S5) and placed onto kidneys for a 3-hour incubation at room temperature with gentle rocking. Kidneys were then washed 3X with TSP for 15 minutes each, and placed into PBS and covered with aluminum foil for storage. The day prior to imaging on a confocal microscope, the tissue was cleared in Refractive Index Matching Solution (RIMS; (40g Histodenz (Sigma D2158) / 0.02M Phosphate Buffer / 0.01% Sodium Azide / 0.1% Tween20 / 1g DABCO) for 3-5 hours, after which the solution was replaced with fresh RIMS. Tissue was then placed into a 35 mm Dish, (No. 1.5 Uncoated Coverslip, 20 mm Glass Diameter MatTek Corporation, P35G-1.5-20-C.S) and covered with enough RIMS to avoid an air bubble. Images were captured using a Nikon A1 Inverted confocal microscope.

### RNA extraction

Kidneys were isolated from e 15.5 mice and placed into ice cold PBS. DNA was collected from tails of embryos for genotyping purposes. RNA was extracted using the Invitrogen PureLink ™ RNA Mini Kit (Thermo Fisher Scientific, 12183025) following the manufacturer protocol. Samples were then sent to Novogene Corporation INC for RNA sequencing.

### Cell type assignment and Deconvolution of bulk RNA

To assign cell types manually we used to VLOOKUP table provided in Table S4 of (Combes et al., 2019) and entered our differentially expressed, filtered bulk gene list (FDR >0.05, −0.5>LogFC>0.5). After all the expression values in NPC were computed by the excel VLOOKUP script, we sorted by imputed cell type, and annotated the gene based on the cell type with unique or highest expression value if LogFC was >0.65. To estimate cell type proportions in bulk RNA-Sequencing data while considering their expression levels in the bulk dataset and the reference data, we performed reference-based decomposition with the R package BisqueMarker (Jew et al., 2020). Briefly, expressed genes with zero variance were filtered out and the remaining expressed gene counts were converted to counts per million. Markers for each cell type from single cell dataset are filtered at FDR < 0.5 and then are ranked based on p-value (Combes et al., 2019). To estimate cell type proportions in bulk samples, Bisque applies a weighted principal component analysis (PCA) approach, where 1st principal component is calculated on the expression matrix using a subset of the marker genes. Expression matrix of markers are scaled and each gene column is multiplied by its weight (log-fold change). We chose the Combes dataset over our own (Jarmas et al., 2021) since at the time of this analysis our data were yet to complete peer review, and the Combes dataset included all cell types whereas ours was cortically biased.

### Quantification

Quantification of distal and proximal tubule markers were done using NIS Elements with help from Dr. Matt Kofron. Binaries were created to measure the total kidney area, and the areas containing distal markers, proximal markers, or GFP (in um^2^). The binaries were kept the same for each slide that was imaged, and only modified to account for GFP gene dose.

### Renal Panel

Littermates were euthanized between day 60 and day 90 for renal panels. Blood was removed within 5 minutes of being euthanized to avoid clotting, placed into serum separator tubes (Fisher Scientific, 02-675-185) and centrifuged at max speed for 5 minutes. Blood samples were then sent to IDEXX Laboratories Inc. to be tested for renal function. The kidneys were also removed at this time for nephron counts.

### Nephron Counts

HCl maceration of whole kidneys was performed according to (MacKay et al., 1987; Peterson et al., 2019). Kidneys isolated from > P28 animals were minced using a razor blade after removal of capsule, and incubated in the presence of 6N HCL for 90 min. The dissociated kidneys were vigorously pipetted every 30 minutes to further disrupt them. After incubation, 5 volumes of distilled water were added to the samples followed by incubation at 4°C overnight. 100ul of the thoroughly mixed macerate was then pipetted into a cell culture dish and glomeruli were counted in triplicate (three aliquots) for each sample. A single experimenter, blinded to the genotypes of the kidneys being scored, performed all counts, to avoid interobserver variability.

### Flow Cytometry

Briefly, P0 kidneys were isolated from Six2^+/Cre;^ Notch2^+/f^ and Six2^+/Cre;^ Notch2^f/f^ littermates, the capsule removed, and the kidneys digested in 250ul of 1mgl/ml Collagenase D shaking in an Eppendorf Thermomixer R at 1400rpm for 10-15 minutes at 37°C to remove the peripheral nephron progenitor cells. The remnant kidney tissue was removed, and the cell suspension was washed twice with 1% BSA/PBS. The number of GFP-expressing cells was quantified by a Sony SH800S flow cytometer (Sony Biotechnology).

### CRISPR/CAS9 mediated deletion of *NCSTN* in the NHSK iPSC line

CRISPR-Cas9 was used to introduce a mutation in an iPSC line (NHSK) derived from skin cells of a female donor with a normal karyotype. Two guide RNAs (gRNAs) were designed to induce a stop codon in exons 2 of the NCSTN gene (AAACTCTCAACTCTCACTGGCAGCC and CACCGGCTGCCAGTGTGAGTTGAAG) (Figure 6A). The gRNA target sequences were selected according to the on- and off-target scores from http://CRISPOR.org web tool (Haeussler et al., 2016) and cloned into the pX458M-HF vector (modified from the pX458 vector, addgene #48138) to carry an optimized sgRNA scaffold and a high-fidelity eSpCas9(1.1)-2A-GFP (Chen et al., 2013; Slaymaker et al., 2016). The editing activity of the plasmid was validated in 293T cells by the T7E1 assay. A single cell suspension of iPSC cells was prepared using accutase and 1×10e6 cells were nucleofected with 5ug of the plasmid using program CA137. Forty-eight hours post-nucleofection, GFP-positive cells were isolated by FACS and replated at cloning density in hESC media containing 20% <u>K</u>nock<u>o</u>ut^™^ <u>S</u>erum <u>R</u>eplacement (KOSR) (Gibco^™^), 4ng/mL bFGF and 10uM Y27632 (inhibitor of Rho-associated, coiled-coil containing protein kinase; ROCK) in 6 well dishes containing 187,500/well mitomycin C-inactivated CF1 MEFs. After 1-2 weeks, single clones with stereotypical iPSC morphology were manually excised and transferred to mTeSR1/Matrigel culture conditions for genotyping, expansion and cryopreservation. PCR and enzyme digestion identified candidates for correctly targeted clones. DNA from two *NCSTN* deficient clones as well as unmodified control (C20) and parental clone (NHSK) was PCR amplified and sequenced to establish the molecular nature of the mutation. NCSTN-/- (C1) and unmodified control (C20) expanded clones were analyzed by Western blot for *NCSTN* expression with an anti-Nicastrin antibody (N1660, Sigma-Aldrich).

### NOTCH1 activation with EDTA

was performed on NCSTN-/- (C1 and C14) expanded clones and parental NHSK iPSCs as previously described (Ilagan et al., 2011; Rand et al., 2000). Briefly, cells were washed with phosphate buffered saline solution (DPBS; Gibco) and then incubated in Trypsin/EDTA (Invitrogen) for 10 min at 37°C. Following cell detachment, mTeSR media was added to each well and cells were incubated at 37°C for 2h. Cells were then collected and western blot analysis was performed with cleaved Notch1 (Val1744) rabbit monoclonal antibody (Stem Cell Biologies).

### Generation of iPSC-derived 3D kidney organoid

hiPSCs (either C20 or C1) were grown on 6 well plates covered with Matrigel (MG) in defined, feeder-free maintenance medium (mTeSR for human ES and iPS cells, Stem Cell Technologies, cat#85850) as previously described (Low et al., 2019; Yu et al., 2007). When cells reached ~80% confluency (day −1), medium was replaced with mTeSR containing 10mM of Rock inhibitor. Differentiation into 3D kidney organoids was performed according to the Morizane et al. protocol (Morizane and Bonventre, 2017) with some modifications. Briefly, on day 0 of the differentiation protocol, cells were dissociated via Accutase (STEMCELL Technologies) and plated in 24 well plates covered with matrigel at 20,000 cells/per well and grown in basic differentiation media (advanced RPMI with Glutamax) with sequential addition of CHIR (8μM) and noggin (days 0-3), followed by Activin A (days 4-6) and FGF9 (10 ng/mL, days 7-9) as previously described (Morizane and Bonventre, 2017), with daily media changes until day 9. On day 9 of differentiation, cells were dissociated with Accutase followed by aggregation on transwell membranes at an air/liquid interphase as previously described (Takasato et al., 2016). Basic differentiation medium supplemented with CHIR (3 μM) and FGF9 (10 ng/mL) was added to the bottom of the transwell. The cell aggregates were then cultured at 37°C, 5% CO2 with daily media changes for additional 12-14 days. On day 9+2 the medium was changed to the basic differentiation medium supplemented with FGF9 10ng/mL (days 9+2 to 9+4). After that, the organoids were cultured in basic differentiation medium with no additional factors (days 9+5 to 9+12). Total number of days to analysis was 21-24.

### Whole mount immunofluorescence of 3D kidney organoids

Organoids were fixed in transwell plates with 4% PFA at 4°C for 20 minutes as previously described (Takasato et al. 2015). 4% PFA was removed and organoids were washed x3 times with Dulbecco’s Phosphate Buffered Saline (DPBS, Gibco). 150μl of blocking buffer (10% donkey serum / 0.3% TritonX / DPBS) was added into each well of a 24 well plate. Organoids were cut off the filter and submerged in the blocking buffer at room temperature for 2-3 hours on a rocker. Primary antibodies including were prepared in blocking buffer (0.3% TritonX / 10% Donkey serum / DPBS) with appropriate dilutions (Supplemental Table S5). Blocking buffer was aspirated off the 24 well plate, and 150μl of primary antibodies solution was added into each well. Organoids with primary antibody were incubated at 4°C over-night while shaking and protected from light. Primary antibody solution was removed off each plate and organoids were washed with PBTX (0.3% TritonX / DPBS). Appropriate secondary antibodies (all 1:400 dilution, Supplemental Table S5) were diluted in PBTX and 150 μl was added into each well in the 24 well plate.

Organoids were incubated in secondary antibody solution at 4 °C over-night while shaking and protected from light. Following removal of secondary antibody solution, organoids were either incubated with 20mg/ml DAPI (1:1000 dilution) in DPBS for 3hr and then washed, or washed with DPBS. Slides were mounted with Prolong Gold (Cell Signaling Technology, 9071S) and kept at 4 °C overnight until visualized on a Nikon A1 Inverted confocal microscope.

## Acknowledgements

We would like to thank Dr. Eric Brunskill for encouragement, advice, and for the creation of the NCSTN-deleted iPSC line, Dr. Yueh-Chiang Hu and the Transgenic Animal and Genome Editing Core facility, the Research Flow Cytometry Core, Dr. Chris Mayhew and the Pluripotent Stem cell facility, and the Confocal Imaging Core at Cincinnati Children’s Hospital Medical Center. We thank Drs. Meredith Schuh, Aaron Zorn and Joo-Seop Park for critical comments.

## Competing interests

The authors declare no competing interests

## Funding

This study was supported by funds from the William K. Schubert Endowment to RK, the National Institute of Diabetes and Digestive and Kidney Diseases (NIH DK106225 to R.K.; 1F30 DK 123841 to A.J.P) as well as generous financial support for the cores at Cincinnati Children’s Hospital Medical Center. The Research Flow Cytometry Core is supported by NIH S10OD023410.

## Data availability

The scRNA-sequencing data analyzed in this study is published in (Jarmas et al., 2021) and deposited in NCBI’s Gene Expression Omnibus (GEO) database under accession codes GSE173264 (https://www.ncbi.nlm.nih.gov/geo/query/acc.cgi?acc=GSE173264), GSE173265 (https://www.ncbi.nlm.nih.gov/geo/query/acc.cgi?acc=GSE173265), and GSE173266 https://www.ncbi.nlm.nih.gov/geo/query/acc.cgi?acc=GSE173266), and contain all processed bam files, raw counts of genes across barcodes/cells and cell annotations in the form of a metafile.

## Notes

### Competing Interest Statement

The authors have declared no competing interest.

